# Knowledge-based antibody repertoire simulation, a novel allele detection tool evaluation and application

**DOI:** 10.1101/2021.07.01.450681

**Authors:** Xiujia Yang, Yan Zhu, Huikun Zeng, Sen Chen, Junjie Guan, Qilong Wang, Chunhong Lan, Deqiang Sun, Xueqing Yu, Zhenhai Zhang

## Abstract

Detailed knowledge of the diverse immunoglobulin germline genes is critical for the study of humoral immunity. Hundreds of alleles have been discovered by analyzing antibody repertoire sequencing (Rep-seq or Ig-seq) data via multiple novel allele detection tools (NADTs). However, the performance of these NADTs through antibody sequences with intrinsic somatic hypermutations (SHMs) is unclear. Here, we developed a tool to simulate repertoires by integrating the full spectrum features of an antibody repertoire such as germline gene usage, junctional modification, position-specific SHM and clonal expansion based on 2152 high-quality datasets. We then systematically evaluated these NADTs using both simulated and genuine Ig-seq datasets. Finally, we applied these NADTs to 687 Ig-seq datasets and identified 43 novel alleles using defined criteria. Twenty-five alleles were validated through findings of other sources. In addition to the novel alleles detected, our simulation tool, the results of our comparison, and the streamline of this process may benefit further humoral immunity studies via Ig-seq.

## Introduction

Genetic variations of antibody germline genes plays a pivotal role in humoral immunity. For instance, the allele variants of IGHV1-69 greatly impact the ability to develop broadly neutralizing antibodies (bNAbs) against influenza virus^1^, and modulate IGHV germline gene utilization^2^. In addition, the polymorphism in IGHV4-61 is associated with a risk in rheumatic heart disease^3^. In fundamental research, accurately assigning germline genes to antibody sequences is also critical. It affects the analysis of clonotype, somatic hypermutation (SHM), and the maturation pathway of antibody clones. Therefore, germline alleles are essential for delineating the ontogeny and evolution of antibody responses specific to antigens or vaccines. Despite this need, a comprehensive collection of novel alleles has not yet been achieved^4^.

The advent of antibody repertoire sequencing (Rep-seq or Ig-seq) technology allows the acquisition of millions of antibody sequences and these unprecedented data facilitate the discovery of novel alleles through tools with specific aims (i.e. novel allele detection tools, NADTs)^5–9^. As antibody sequences undergo extensive SHMs along with B cell proliferation once activated by an antigen, novel allele detection for antibody genes are more challenging than traditional mutation detection in conventional genes where only base errors caused by PCR and high-throughput sequencing (HTS) need to be considered^6^. To distinguish SHMs and base errors from real polymorphisms, NADTs use distinct algorithms and are supposed to be effective in typical scenarios.

Algorithm wise, *TIgGER*^6^, *LymAnalyzer*^8^, and *Partis*^7^ employ a SNP-based approach. Novel alleles are predicted by identifying SNPs in the reference germlines. For example, *TIgGER* and *Partis* employ mutation accumulation plots to identify SNPs. Therefore, the major challenge for these NADTs is to distinguish SNPs from SHMs. In contrast, *IgDiscover*^5^ annotates the input sequences with an initial germline database to form clusters and subsequently predicts novel alleles based on consensus building within clusters. This sequence-based approach circumvents the SNP set determination procedure encountered by the SNP-based approach and can easily output the novel germline sequences regardless of the distances to their nearest counterparts. Nevertheless, it heavily relies on repertoire types and is suggested to work efficiently only on naïve repertoires featured by a substantial fraction of unmutated sequences. *IMPre*^9^ uses a seed-based approach. It starts with a seed sequence and extends the sequence in both directions if defined requirements are met. It is worth mentioning that both the sequence-based approach and the seed-based extension approach can identify novel alleles that have insertions and deletions compared to the known germlines.

Despite these algorithm differences, it remains unclear how NADTs above compete with each other in practice. A previous study presented a comparison among 3 NADTs (i.e. *IgDiscover*, *TIgGER* and *Partis*)^7^, but the study was not comprehensive as to both the number of included NADTs and the kind of challenges that need to be overcome in novel allele detection. To evaluate the five NADTs *TIgGER*, *LymAnalyzer*, *Partis, IgDiscover* and *IMPre* objectively, we used a repertoire simulation tool that incorporates the full spectrum of repertoire features extrapolated from 2152 datasets, including germline gene usage, junctional modification, position-specific SHM and clonal expansion. We then systematically evaluated these NADTs using both the simulated datasets and paired genuine bulk and single-cell repertoire sequencing datasets. We identified 43 alleles from 683 datasets using the criterion set based on the comparison result. This systematic evaluation, together with the novel alleles we present here, may aid future novel allele identification and thus achieve a better interpretation of adaptive immune receptor repertoire sequencing (AIRR-seq) dataset.

## Results

### An overview of 5 NADTs and the study design

To perform solid and comprehensive comparison for currently available NADTs, we employed *TIgGER*^6^, *IMPre*^9^, *IgDiscover*^5^, *LymAnalyzer*^8^ and *Partis*^7^. Their basic information is summarized in Table 1. As these five NADTs were developed using various programming languages, their installations are subject to various dependencies. With respect to their applications, *IMPre* and *LymAnalyzer* work on both T cell receptor (TCR) and B cell receptor (BCR) while the other three only work on BCR. All NADTs support both heavy chain (IGH) and light chain (IGK and IGL) of BCR, while *IMPre* and *LymAnalyzer* also support TRB and TRA. *TIgGER* and *Partis* only support V genes, *IMPre* and *LymAnalyzer* support V and J genes, while *IgDiscover* supports V, D, and J genes. Except *IgDiscover* and *LymAnalyzer*, all other NADTs underwent *in silico* benchmark during development. *Partis* developers compared their NADT with others, but no systematic third-party comparison has been performed among them. Therefore, a comprehensive and systematic comparison would benefit the field for novel allele detection using antibody repertoire datasets.

**Table 1.**
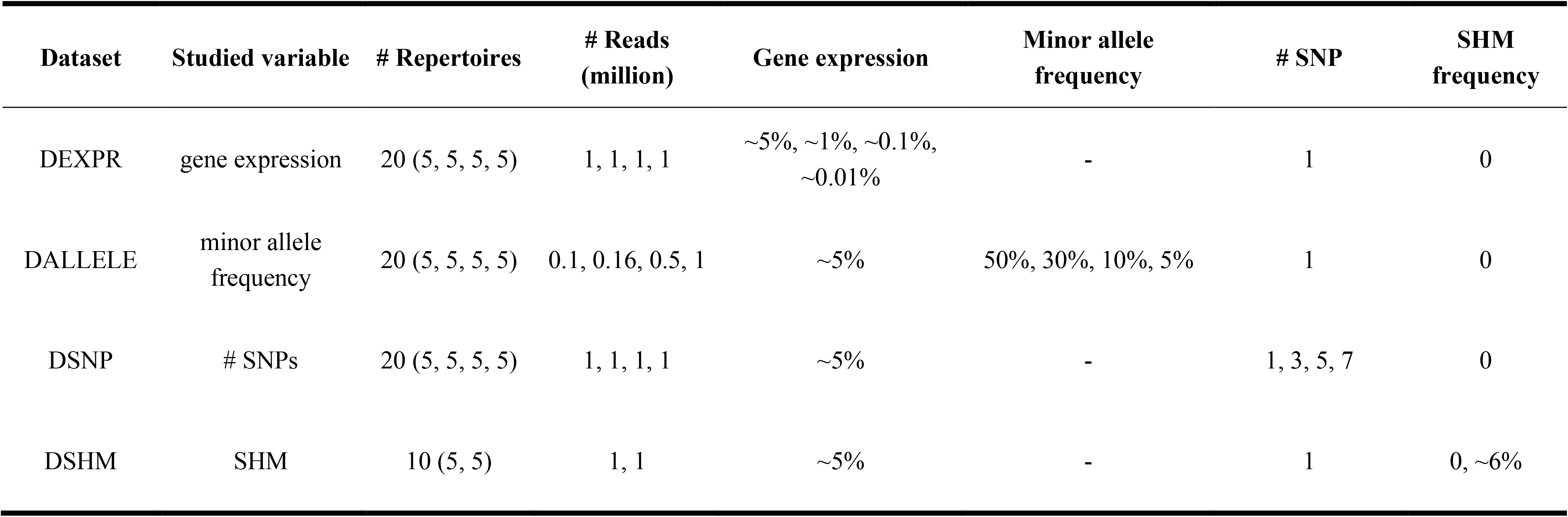
Characterization of four simulated datasets.

| Dataset | Studied variable | # Repertoires | # Reads<br>(million) | Gene expression | Minor allele<br>frequency | # SNP | SHM<br>frequency |
| --- | --- | --- | --- | --- | --- | --- | --- |
| DEXPR | gene expression | 20 (5, 5, 5, 5) | 1, 1, 1, 1 | ~5%, ~1%, ~0.1%,<br>~0.01% | - | 1 | 0 |
| DALLELE | minor allele<br>frequency | 20 (5, 5, 5, 5) | 0.1, 0.16, 0.5, 1 | ~5% | 50%, 30%, 10%, 5% | 1 | 0 |
| DSNP | # SNPs | 20 (5, 5, 5, 5) | 1, 1, 1, 1 | ~5% | - | 1, 3, 5, 7 | 0 |
| DSHM | SHM | 10 (5, 5) | 1, 1 | ~5% | - | 1 | 0, ~6% |

When we compared the supportive features of these NADTs, we found *IMPre* to be the most versatile and user-friendly NADT before considering its performance for novel allele detection (Supplementary Table 1). To gain more insights into these NADTs, we evaluated their performance with both simulated and real-world Ig-seq datasets (Figure 1) The benchmark result was then summarized and translated into knowledge-based filtration criteria used to obtain credible novel alleles from collected bulk sequencing dataset.

**Figure 1.**
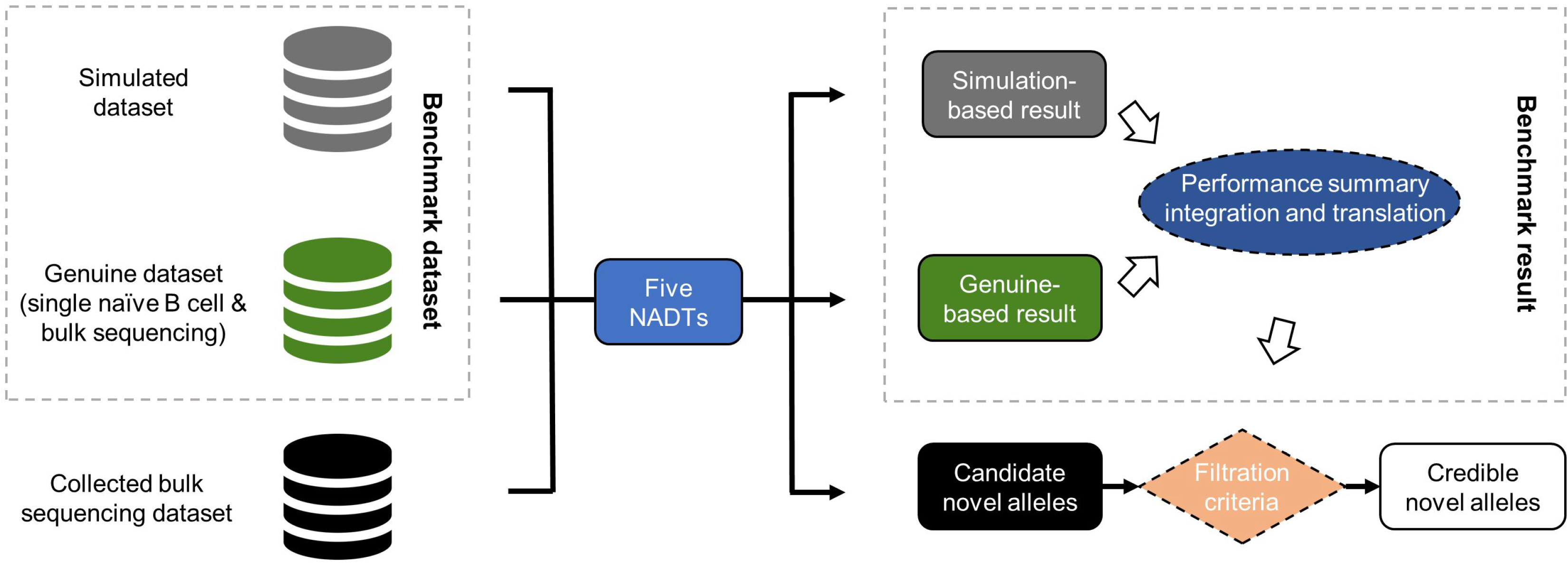
Schematic overview of the study design. In this study, both in silico simulated and genuine Ig-seq dataset were employed as benchmark datasets that serve as the input of all five NADTs independently. The performances of these NADTs were then summarized and integrated, and translated into filtration criteria capable of facilitating the evaluation of candidate novel alleles. Among all candidate novel alleles reported based on the collected bulk sequencing dataset, we retained only those credible novel alleles passing the defined filtration criteria.

### A flexible immune repertoire sequencing dataset simulation tool and the benchmark dataset

Generating *in silico* Ig-seq datasets is a challenging task. An ideal Ig-seq simulating tool should reflect the preferential gene usage, junctional nucleotide insertion and deletion, phylogenetic clonal structure, various allele ratio, and the base errors intrinsic to PCR amplification and next-generation sequencing (NGS). Although several repertoire simulation tools exist^7,10–13^, none of them incorporate the full features of Ig-seq dataset. Therefore, we built *IMPlAntS* (**I**ntegrated and **M**odular **P**ipe**l**ine for **Ant**ibody Repertoire **S**imulation) and it enables both one-stop repertoire simulation and modular calls for adaption to customized pipelines.

Briefly, *IMPlAntS* consists of three consecutive steps, i) generation of independent V(D)J rearrangements; ii) generation of SHM with phylogenetic structure within clones; and iii) generation of NGS reads incorporating base errors (Supplementary Figure 1). These steps can be implemented individually or collectively using the corresponding scripts.

In the first step, a series of key parameters can be specified in the configuration files. These parameters include V(D)J gene usage, allele ratio, the distribution of insertion and deletion length, and the percentage of productive rearrangements. In the second step, we generated SHMs in rearranged sequences in a way similar to that reported by Yermanos et al.^13^ to create the phylogenetic sequences as in the real repertoire. The resultant repertoire with SHMs comes from several iterations of introducing SHMs to the selected sequences based on the positional mutability and substitutability models. These two models, together with the parameters involved in the first step, derive from our previous large-scale study^14^. Finally, we employed a popular NGS simulation tool, ART, to produce NGS reads^15^. More details for *IMPlAnts* can be found on github (https://github.com/Xiujia-Yang/IMPlAntS).

With this pipeline, we generated four datasets: DEXPR, DSNP, DALLELE, and DSHM (Table 2). Noteworthy is that only DSHM was generated with all three steps mentioned above. In contrast, the other three datasets were generated with only the first and the final step as they contain no SHMs. Each of the four datasets was comprised of 20 repertoires, except for DSHM (n=10). The constituent repertoires within each dataset contained variation only in the studied variable. Except for DSHM, each dataset contained four groups (two groups for DSHM). While each group is represented by five repertoire replicates and has a distinct level as to the studied variables. Other variables were set identically among groups within each dataset and to a level theoretically most favorable to novel allele detection. For each repertoire, we generated 1 million reads to avoid the read number limitation mentioned in the *IgDiscover* manual (at least 750,000 was recommended). The only exception was with DALLELE, in which repertoires in different groups had varying numbers of reads to make the novel alleles represented by the same number of reads. Lastly, we artificially created “novel” alleles by random selection of the positions and SNPs in germline sequences. The resultant “novel” alleles together with known ones then served as the initial germline database for NADTs’ benchmarking (Materials and Methods).

**Table 2.**
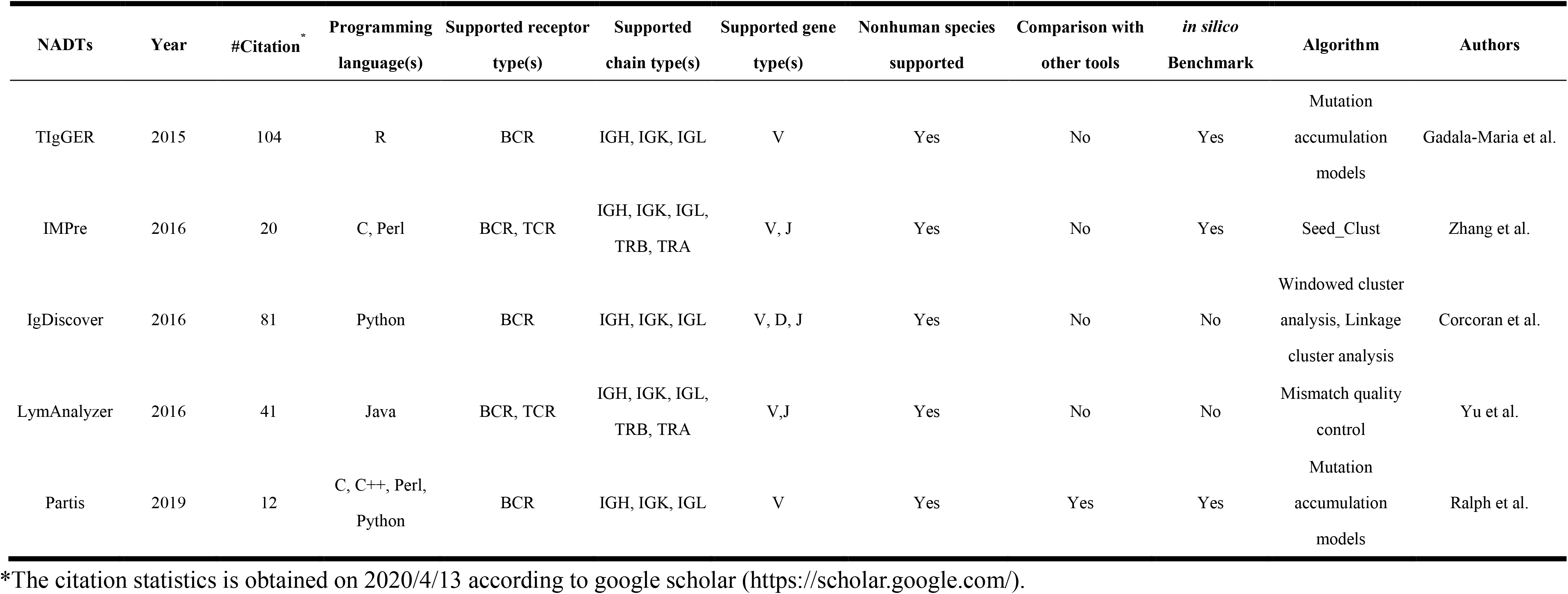
The basic information for 5 NADTs.

| NADTs | Year | #Citation<sup>*</sup> | Programming language(s) | Supported receptor type(s) | Supported chain type(s) | Supported gene type(s) | Nonhuman species supported | Comparison with other tools | *in silico* Benchmark | Algorithm | Authors |
| --- | --- | --- | --- | --- | --- | --- | --- | --- | --- | --- | --- |
| TIgGER | 2015 | 104 | R | BCR | IGH, IGK, IGL | V | Yes | No | Yes | Mutation accumulation models | Gadala-Maria et al. |
| IMPre | 2016 | 20 | C, Perl | BCR, TCR | IGH, IGK, IGL, TRB, TRA | V, J | Yes | No | Yes | Seed_Clust | Zhang et al. |
| IgDiscover | 2016 | 81 | Python | BCR | IGH, IGK, IGL | V, D, J | Yes | No | No | Windowed cluster analysis, Linkage cluster analysis | Corcoran et al. |
| LymAnalyzer | 2016 | 41 | Java | BCR, TCR | IGH, IGK, IGL, TRB, TRA | V,J | Yes | No | No | Mismatch quality control | Yu et al. |
| Partis | 2019 | 12 | C, C++, Perl, Python | BCR | IGH, IGK, IGL | V | Yes | Yes | Yes | Mutation accumulation models | Ralph et al. |

\*The citation statistics is obtained on 2020/4/13 according to google scholar (https://scholar.google.com/).

### Evaluation of the 5 NADTs using *in silico* simulated benchmark dataset

To compare the sensitivity and specificity of the 5 NADTs in detecting novel alleles (allele level) (*LymAnalyzer* was excluded as it reported only SNPs) as well as SNPs (SNP level) (Materials and Methods), we used our *in silico* simulated datasets (Table 3). As expected, lower gene or allele expression and more SNP or SHMs hampered both sensitivities and specificities for at least one NADT in the detection of novel alleles and SNPs in general (Table 3 and Supplementary Figure 2). We found *TIgGER* to work well with respect to both sensitivities and specificities with DEXPR and DSNP, although it did not identify alleles in DALLELE. *IMPre*, though exhibting lower sensitivities and specificities, identified novel alleles in the datasets with all four variables. *IgDiscover* manifested very good specificities although it identified fewer alleles than *TIgGER*. The performance of *Partis* was less optimal in DSNP than that of *TIgGER* but excelled in DALLELE and higher SHM datasets. As *LymAnalyzer* only reports SNPs, it was excluded from allele level comparisons. However, it also showed high sensitivities in all situations in SNP level although the sensitivities were less ideal. The performance of other NADTs was similar in SNP level to that of the allele level.

**Table 3.** Sensitivity and specificity of novel allele detection for 5 NADTs based on four simulated datasets.

| Type | Measurement | Tool | Dataset |  |  |  |  |  |  |  |  |  |  |  |  |  |
| --- | --- | --- | --- | --- | --- | --- | --- | --- | --- | --- | --- | --- | --- | --- | --- | --- |
|  |  |  | DEXPR |  |  |  | DALLELE |  |  |  | DSNP |  |  |  | DSHM |  |
|  |  |  | ~5% | ~1% | ~0.1% | ~0.01% | 50% | 30% | 10% | 5% | 1 | 3 | 5 | 7 | 0% | 6% |
| Allele level | Sensitivity | TIgGER | 1.00 | 0.76 | 0.00 | 0.00 | 0.00 | 0.00 | 0.00 | 0.00 | 1.00 | 0.96 | 1.00 | 0.96 | 1.00 | 0.00 |
|  |  | IMPre | 0.28 | 0.92 | 0.44 | 0.00 | 0.52 | 0.56 | 0.40 | 0.52 | 0.40 | 0.44 | 0.16 | 0.28 | 0.20 | 0.52 |
|  |  | IgDiscover | 0.60 | 0.00 | 0.00 | 0.00 | 0.00 | 0.00 | 0.00 | 0.00 | 0.72 | 0.68 | 0.88 | 0.64 | 0.80 | 0.00 |
|  |  | LymAnalyzer | - | - | - | - | - | - | - | - | - | - | - | - | - | - |
|  |  | Partis | 1.00 | 1.00 | 0.32 | 0.00 | 0.48 | 0.20 | 0.20 | 0.20 | 1.00 | 0.28 | 0.04 | 0.00 | 1.00 | 0.92 |
|  | Specificity | TIgGER | 1.00 | 1.00 | 0.00 | 0.00 | 0.00 | 0.00 | 0.00 | 0.00 | 1.00 | 0.96 | 1.00 | 0.75 | 1.00 | 0.00 |
|  |  | IMPre | 0.17 | 0.44 | 0.24 | 0.00 | 0.63 | 0.70 | 0.33 | 0.28 | 0.25 | 0.30 | 0.09 | 0.21 | 0.13 | 0.56 |
|  |  | IgDiscover | 1.00 | 0.00 | 0.00 | 0.00 | 0.00 | 0.00 | 0.00 | 0.00 | 1.00 | 1.00 | 1.00 | 0.97 | 1.00 | 0.00 |
|  |  | LymAnalyzer | - | - | - | - | - | - | - | - | - | - | - | - | - | - |
|  |  | Partis | 1.00 | 1.00 | 0.80 | 0.00 | 0.82 | 0.90 | 0.90 | 1.00 | 0.97 | 0.30 | 0.05 | 0.00 | 1.00 | 0.67 |
| SNP level | Sensitivity | TIgGER | 1.00 | 0.76 | 0.00 | 0.00 | 0.00 | 0.00 | 0.00 | 0.00 | 1.00 | 0.99 | 1.00 | 0.99 | 1.00 | 0.00 |
|  |  | IMPre | 0.80 | 0.92 | 0.44 | 0.00 | 0.60 | 0.56 | 0.40 | 0.52 | 0.76 | 0.73 | 0.74 | 0.67 | 0.76 | 0.56 |
|  |  | IgDiscover | 0.60 | 0.00 | 0.00 | 0.00 | 0.00 | 0.00 | 0.00 | 0.00 | 0.72 | 0.68 | 0.88 | 0.64 | 0.80 | 0.00 |
|  |  | LymAnalyzer | 1.00 | 1.00 | 1.00 | 0.00 | 1.00 | 1.00 | 1.00 | 1.00 | 1.00 | 1.00 | 1.00 | 1.00 | 1.00 | 1.00 |
|  |  | Partis | 1.00 | 1.00 | 0.32 | 0.00 | 0.48 | 0.20 | 0.20 | 0.20 | 1.00 | 0.44 | 0.30 | 0.04 | 1.00 | 0.92 |
|  | Specificity | TIgGER | 1.00 | 1.00 | 0.00 | 0.00 | 0.00 | 0.00 | 0.00 | 0.00 | 1.00 | 1.00 | 1.00 | 0.70 | 1.00 | 0.00 |
|  |  | IMPre | 0.23 | 0.31 | 0.15 | 0.00 | 0.40 | 0.39 | 0.16 | 0.17 | 0.31 | 0.59 | 0.64 | 0.75 | 0.25 | 0.45 |
|  |  | IgDiscover | 1.00 | 0.00 | 0.00 | 0.00 | 0.00 | 0.00 | 0.00 | 0.00 | 1.00 | 1.00 | 1.00 | 0.94 | 1.00 | 0.00 |
|  |  | LymAnalyzer | 0.09 | 0.09 | 0.09 | 0.00 | 0.11 | 0.10 | 0.09 | 0.08 | 0.09 | 0.23 | 0.34 | 0.31 | 0.09 | 0.00 |
|  |  | Partis | 1.00 | 1.00 | 0.80 | 0.00 | 0.78 | 0.85 | 0.85 | 1.00 | 0.97 | 0.78 | 0.68 | 0.10 | 1.00 | 0.39 |

Taken together, *TIgGER*, *IgDiscover*, and *Partis* showed comparably high specificities and therefore the alleles identified were more reliable. *IMPre* and *LymAnalyzer* provided more allele candidates, but none of the NADTs performed well in all situations. However, each of these datasets was simulated with only one variable with a particular quantity to evaluate the effect of these quantitative measures on the performance of NADTs whereas real-world repertoires always consist of combinations of all variables in multiple quantitative measures.

### Evaluation of 5 NADTs using a combination of single-cell and bulk sequencing dataset

An ideal situation to test NADTs is to genotype all the V alleles in a genome and then compare them with NADTs’ predictions. However, given the high similarities of V alleles and other interspersed tandem sequences among them, sequencing this peculiar region of the genome alone is a challenging task^16^. Therefore, we took an alternative approach by acquiring germline V allele sequences from single-cell repertoire sequencing of naïve B cells (scRep-seq) and then conducted novel allele identification on the bulk Ig-seq datasets from the same donor. The naïve state of antibody sequences and the super-high depth of the scRep-seq data ensured the accuracy of acquired germline sequences. Thus, this evaluation represents the real-world situation.

With the single naïve B cell sequencing dataset, we identified 4 unique novel alleles from 3 donors using a customized pipeline (Table 4, Materials and Methods). All identified novel alleles are minor alleles of the involved genes, with expression ratios to the major ones ranging from 0.19 to 0.89. Moreover, each of them only harbors one SNP compared to their nearest known alleles.

**Table 4.** Novel alleles identified based on single naïve B cell sequencing dataset from 3 donors.

| Nearest known allele<sup>a</sup> | Known allele | # Supportive contigs<sup>b</sup> | Length (bp) | Start | End | SNP loci<sup>c</sup> | Individual |
| --- | --- | --- | --- | --- | --- | --- | --- |
| IGHV7-4-1*02 | IGHV7-4-1*02 | 44 (136, 0.32) | 296 | 1 | 296 | G92A | Donor1 |
| IGHV3-30*18<sup>T</sup> | IGHV3-30*18 | 126 (492, 0.26) | 296 | 1 | 296 | C72G | Donor2 |
| IGHV3-7*03<sup>T</sup> | IGHV3-7*03 | 96 (108, 0.89) | 296 | 1 | 296 | G46A | Donor3 |
| IGHV3-53*04<sup>T,G</sup> | IGHV3-53*01 | 24 (126, 0.19) | 293 | 1 | 293 | T261C |  |

**Note:** **a**, The novel alleles identified by TIgGER using bulk sequencing of IgM sequences are marked with "T" while IgDiscover with "G". **b**, The numbers in the parentheses denote the number of contigs supportive of its known germline variant in the second column and the ratio of the two germline variants. **c**, The indexes in SNP loci are 1-based. Novel allele highlighted in red is not included in the collected germline sequences (see Supplementary Table 7).

We then applied NADTs to the bulk sequencing datasets and compared their novel allele predictions. *TIgGER* identified three and *IgDiscover* identified one out of the four novel alleles (Table 4) while *IMPre* and *Partis* missed all of them. Although *LymAnalyzer* identified two positive SNPs from two novel alleles, it also falsely predicted 14 and 6 SNPs in these two alleles, respectively. In addition, we found two possible novel germline sequences that harbor a considerable number of mismatches with their nearest known germline sequences (Supplementary Table 2). Notably, the novel germline sequence nearest to IGHV1-NL1*01 was identified in 2 of 3 enrolled donors.

Because of the limited number of *bona fide* novel alleles, this evaluation was less comprehensive. We thus exploited the genuine Rep-seq dataset (donor1 and donor3) in another way. As many germline sequences were known through the single naïve B cell sequencing dataset, we artificially generated “novel” alleles as those in the simulated dataset mentioned above and evaluated these NADTs in the same way (Materials and Methods). However, because there were not enough genes expressing at around 0.01%, we did not generate novel alleles at this level. Moreover, the allele ratio is hard to precisely infer even with the single naïve B cell sequencing dataset and was thus left unstudied. We denoted the genuine dataset with different initial databases as GD-EXPR, GD-SNP, and GD-SHM.

The genuine dataset-based benchmark result exhibited a similar performance spectrum as that based on the simulated dataset (Table 5). These similarities included, **i)** each of the three studied factors was found to be influential for at least one NADT for novel allele detection (Supplementary Figure 4), **ii)** *TIgGER* and *IgDiscover* were superior to *IMPre* and *Partis* in both sensitivity and specificity for detecting novel alleles with multiple SNPs (i. e. 3, 5, and 7) (Supplementary Figure 5), **iii)** the SNP-level performance spectrum in general resembled that of allele level, **iv)** *IMPre* and *Partis* presented higher sensitivity and specificity for identifying SNPs in DSNP than for alleles, and **v)** *LymAnalyzer* remained the most sensitive but least specific NADT in identifying SNPs.

**Table 5.** Sensitivity and specificity of novel allele detection for 5 NADTs based on genuine Ig-seq dataset.

| Type | Measurement | Tool | Dataset |  |  |  |  |  |  |  |  |
| --- | --- | --- | --- | --- | --- | --- | --- | --- | --- | --- | --- |
|  |  |  | GD-EXPR |  |  | GD-SNP |  |  |  | GD-SHM |  |
|  |  |  | ~5% | ~1% | ~0.1% | 1 | 3 | 5 | 7 | IgM | IgG |
| Allele level | Sensitivity | TIgGER | 0.80 | 0.60 | 0.20 | 0.80 | 0.76 | 0.80 | 0.60 | 0.80 | 0.00 |
|  |  | IMPre | 0.40 | 0.64 | 0.04 | 0.40 | 0.36 | 0.08 | 0.40 | 0.40 | 0.00 |
|  |  | IgDiscover | 1.00 | 1.00 | 0.40 | 1.00 | 1.00 | 1.00 | 0.84 | 1.00 | 0.80 |
|  |  | LymAnalyzer | - | - | - | - | - | - | - | - | - |
|  |  | Partis | 0.56 | 0.48 | 0.00 | 0.60 | 0.00 | 0.00 | 0.00 | 0.60 | 0.32 |
|  | Specificity | TIgGER | 0.57 | 0.75 | 0.14 | 0.57 | 0.41 | 0.52 | 0.31 | 0.57 | 0.00 |
|  |  | IMPre | 0.27 | 0.33 | 0.03 | 0.28 | 0.21 | 0.04 | 0.23 | 0.27 | 0.00 |
|  |  | IgDiscover | 1.00 | 1.00 | 0.33 | 1.00 | 0.63 | 0.81 | 0.70 | 1.00 | 1.00 |
|  |  | LymAnalyzer | - | - | - | - | - | - | - | - | - |
|  |  | Partis | 0.17 | 0.12 | 0.00 | 0.18 | 0.00 | 0.00 | 0.00 | 0.17 | 0.43 |
| SNP level | Sensitivity | TIgGER | 0.80 | 0.60 | 0.40 | 0.80 | 0.79 | 0.80 | 0.63 | 0.80 | 0.00 |
|  |  | IMPre | 0.40 | 0.76 | 0.04 | 0.48 | 0.71 | 0.18 | 0.73 | 0.40 | 0.00 |
|  |  | IgDiscover | 1.00 | 1.00 | 0.60 | 1.00 | 1.00 | 1.00 | 0.84 | 1.00 | 0.80 |
|  |  | LymAnalyzer | 1.00 | 1.00 | 0.72 | 1.00 | 1.00 | 1.00 | 1.00 | 1.00 | 0.96 |
|  |  | Partis | 0.56 | 0.48 | 0.00 | 0.60 | 0.17 | 0.11 | 0.07 | 0.60 | 0.32 |
|  | Specificity | TIgGER | 0.57 | 0.75 | 0.18 | 0.57 | 0.49 | 0.67 | 0.53 | 0.57 | 0.00 |
|  |  | IMPre | 0.07 | 0.03 | 0.00 | 0.02 | 0.10 | 0.02 | 0.12 | 0.05 | 0.00 |
|  |  | IgDiscover | 1.00 | 1.00 | 0.43 | 1.00 | 0.60 | 0.86 | 0.79 | 1.00 | 1.00 |
|  |  | LymAnalyzer | 0.00 | 0.00 | 0.00 | 0.00 | 0.01 | 0.01 | 0.02 | 0.00 | 0.00 |
|  |  | Partis | 0.08 | 0.08 | 0.00 | 0.09 | 0.08 | 0.11 | 0.07 | 0.08 | 0.34 |

However, several variations were also remarkable and they included, **i)** for genuine datasets, *TIgGER* and *IgDiscover* performed better in identifying novel alleles expressed at a low level (i. e. ~0.1%) than for simulated dataset; both were thus superior to *IMPre* and *Partis*,**ii)** although *Partis* remained excellent in overcoming SHM noise, it was outperformed by *IgDiscover* (Supplementary Figure 5), which exhibited a surprisingly high sensitivity of 0.80 and specificity of 1.00 at both SNP and allele levels, **iii)** *IgDiscover* manifested significantly higher sensitivities and specificities than *TIgGER* in three datasets, and **iv)** *LymAnzlyzer* displayed low and negligible specificities.

To seek the underlying reasons accounting for these discrepancies, we assessed the output of these NADTs as well as the properties of each input dataset. We found that the inferior performance of *IgDiscover* and *TIgGER* on DEXPR in the simulated dataset was caused by low sequence identities to the germline (Supplementary Figure 6), which was caused by sequencing errors simulated by ART, the NGS simulator. Similarly, the failure in DSHM for *IgDiscover* could also be attributed to the paucity of unmutated sequences as a consequence of the simulation of SHMs and NGS errors (Supplementary Figure 7). In contrast, the success in GD-SHM for *IgDiscover* indicated that a number of unmutated sequences also exist in IgG dataset. We therefore determined the frequency of such sequences for each simulated novel allele and found that it ranged from 0.07% to 1.11%, with a median of 0.31% (Supplementary Figure 7), which agrees with reported values of previous studies^17,18^. Interestingly, *TIgGER* failed to detect any novel alleles from both simulated and genuine datasets containing SHMs as its algorithm is expected to be more robust to datasets with SHMs. Our in-depth analyses showed that *TIgGER* failed to identify novel alleles for DSHM and GD-SHM for different reasons. For DSHM, the simulated SHMs created an overly-diversified repertoire, in which plural sequences for each novel allele were too rare to pass the threshold *min_seqs*. Whereas in GD-SHM, the diversity of sequences perfectly matching novel alleles failed to meet the default threshold *j_max*. In addition, we noted a remarkable difference in the diversity filtration criterion between *TIgGER* and *IgDiscover:TIgGER* employs a quantitative filtration (*j_max*) whereas *IgDiscovers* uses a qualitative filtration (*CDR3_exact*). When considering only the diversity criterion, *TIgGER* is more strict than *IgDiscover,* and this explains the compromised performance of *TIgGER*. Finally, the lower specificity of *LymAnalyzer* in the genuine dataset may account for the non-independent mutation events in a genuine dataset that tends to be interpreted as SNPs according to its algorithm.

Together, we concluded that **i)** *TIgGER* and *IgDiscover* outperform all other NADTs considering both sensitivity and specificity in most situations, **ii)** *Partis* is characterized by remarkable robustness in overcoming the challenge imposed by SHMs, **iii)** *IMPre* is outstanding in detecting minor alleles, and **iv)** *LymAnalyzer* is sensitive at the cost of specificity.

### Forty-three novel alleles are identified from a total number of 687 Ig-seq datasets

With the knowledge obtained above, we designed a scheme to identify reliable novel alleles using 4 NADTs (excluding *LymAnalyzer*) from bulk Ig-seq dataset. As intrinsic features (i.e. expression level, allele ratio, and number of SNPs to the nearest allele) of novel alleles were unknown, we took into account the overall performance of each NADT summarized above and gave more credit to *TIgGER* and *IgDiscover.* We classified all Ig-seq datasets into two groups with regard to the SHM richness according to the isotypes (Material and Methods). For IgM datasets, novel alleles found by at least 2 NADTs with at least one being either *TIgGER* or *IgDiscover* were retained. For datasets in which SHMs were expected to be enriched, only alleles called by two out of three NADTs, namely *TIgGER*, *IgDiscover* and *Partis*, were retained.

We then explored the efficiency of this scheme in identifying novel alleles from a total number of 424 Ig-seq datasets either generated in-house or from the public resource (Material and Methods). The selected datasets stemmed from 382 donors and were all obtained from RNA samples amplified with RACE (rapid amplification of cDNA ends) protocols. Detailed metadata for these datasets are outlined in Supplementary Table 3. According to the dataset classification criteria (Material and Methods), we obtained 336 (79.2%) SHM-rich datasets (enriched for IgG sequences) and 88 (20.8%) SHM-sparse datasets (enriched for IgM sequences) (Figure 2A). Despite the lower fraction in overall datasets, IgM datasets contain more reads than IgG datasets (Figure 2B). Applying the selected 4 NADTs to these datasets, we found clear differences between the four NADTs in both the number of samples identified with novel alleles and the number of unique novel alleles (Table 6). *IMPre* discerned novel alleles for 71.0% of the datasets, whereas the other three NADTs found novel alleles for only 16.3% to 18.2% datasets. Moreover, the other three NADTs reported novel alleles for a sharply lower (over 10-fold) percentage of SHM-rich datasets than SHM-sparse datasets, which was likely due to more SHMs and low number of input reads that had reduced the confidence for these NADTs to make novel calls. In contrast, *IMPre* reported novel alleles for a large fraction of IgG datasets (63.4%) and also more novel alleles overall for individual samples (Table 6 and Figure 2C), which likely reflects its higher sensitivity to those underrepresented sequences (Table 3). However, the genuine sensitivity and specificity for the NADTs were elusive through these bulk sequencing datasets, for which we have no access to the genotype information.

**Figure 2.**
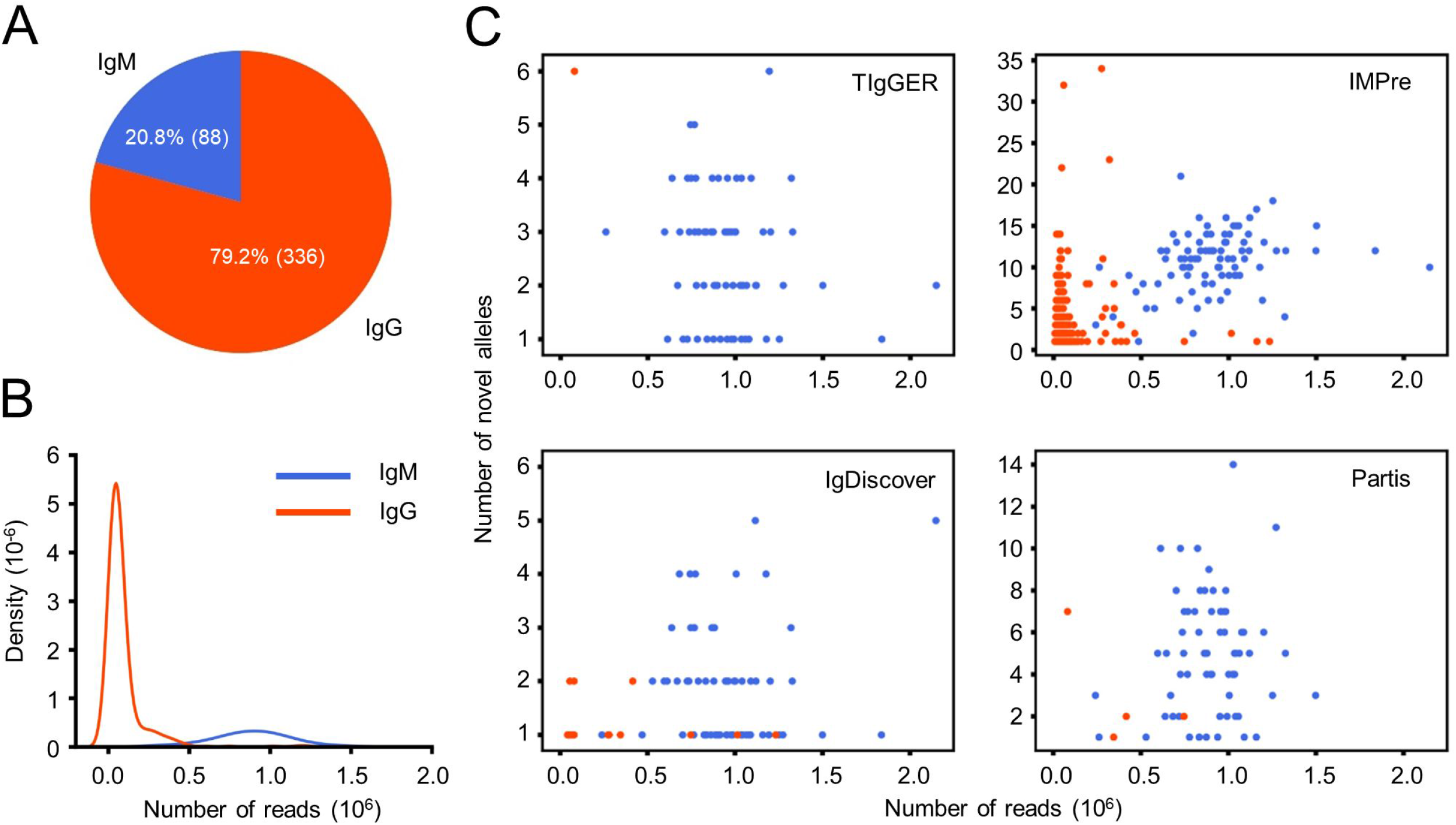
Quantitative characterization of Ig-seq datasets and novel alleles identified by 4 NADTs. **(A)** Composition of IgM (SHM-sparse) and IgG (SHM-enriched) datasets. **(B)** Density of Ig-seq datasets with different number of reads. **(C)**. Correlation between the number of novel alleles for each dataset and the number of reads. Note that only dataset reported with novel alleles by a certain tool is included.

**Table 6.** Quantitative summary of novel alleles identified from Ig-seq datasets by 4 NADTs.

| NADTs | # Datasets (IgG) | # Datasets (IgM) | # Unique novels (IgG) | # Unique novels (IgM) | # Datasets (total) | # Unique novels (total) |
| --- | --- | --- | --- | --- | --- | --- |
| TIgGER | 1 (0.3) | 68 (77.3) | 6 (0.8) | 57 (4.8) | 69 (16.3) | 57 (4.8) |
| IMPre | 213 (63.4) | 88 (100.0) | 740 (96.1) | 1033 (86.5) | 301 (71.0) | 1033 (86.5) |
| IgDiscover | 15 (4.5) | 62 (70.5) | 16 (2.1) | 50 (4.2) | 77 (18.2) | 50 (4.2) |
| Partis | 4 (1.2) | 65 (73.9) | 12 (1.6) | 101 (8.5) | 69 (16.3) | 101 (8.5) |
| **Total** | **215 (64.0)** | **88 (100.0)** | **770** | **1194** | **303 (71.5)** | **1194** |

Note, the number in each parentheses indicates the corresponding percentage (%) of each item. For columns indicating number of datasets, the associated percentages were calculated based on the total number of datasets (or of a specific type, see Figure 2A).

Applying this scheme to 424 Ig-seq datasets, we identified 23 and 2 reliable novel alleles from SHM-sparse and SHM-rich group, respectively (Supplementary Table 4). One novel allele, IGHV3-33*01_G72C, was identified in both groups. Three of the 24 unique novel alleles were found to harbor more than one SNPs to their corresponding nearest alleles, while eleven were found in more than one donor (Figure 3). The most frequent novel allele was found in 29 donors. Notably, 17 of the 24 novel alleles can also be identified from public databases or independent reports in the literature (Materials and Methods) (Supplementary Table 4), which demonstrated the high efficiency of our scheme. To enlarge the knowledge database of novel alleles, we also included 263 multiplex datasets we collected in a previous study into our analysis^14^. These latter datasets were derived from 71 donors and consisted of 186 SHM-rich datasets and 77 SHM-sparse datasets. Considering the degenerate primers designed against framework region 1 (FR1) of V genes, we considered only the sequence downstream of FR1 for each novel allele for these multiplex datasets. Applying the same scheme to these datasets as to RACE datasets, we identified in total 22 novel alleles (Supplementary Table 5) and found that 21 of them were from SHM-sparse datasets. Eleven of the 22 novel alleles can be found in previous public or published resources. Combining the two novel allele sets, we identified 43 unique novel allele sequences from a total number of 687 Ig-seq datasets (3 novel alleles were found in both RACE and multiplex dataset).

**Fig. 3.**
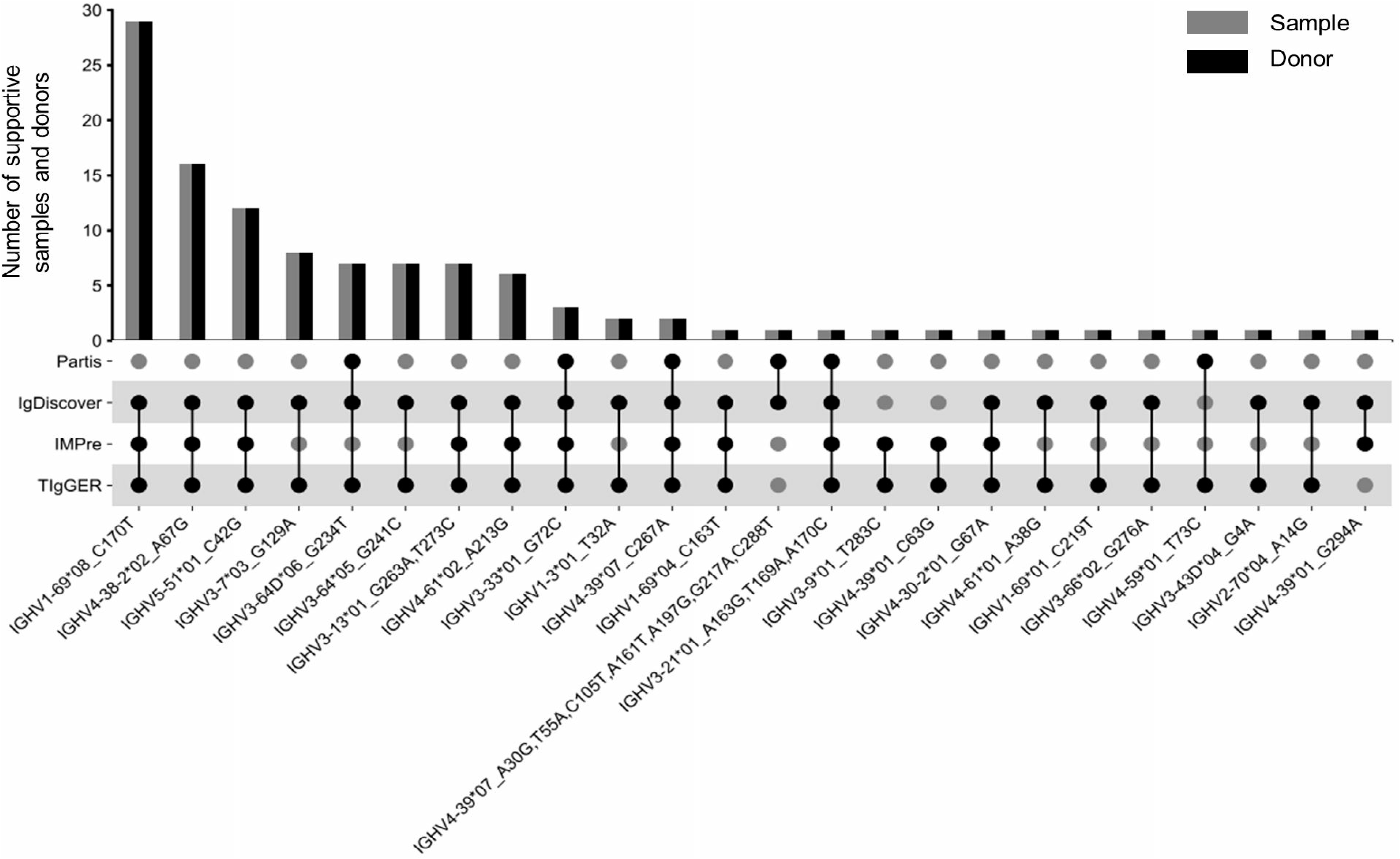
Twenty-four novel alleles identified from 424 Ig-seq datasets amplified using RACE protocol. The top bar graph shows the number of supportive samples and donors. The bottom scatter plot shows the set of tools identifying a typical novel allele. Numbers in the x-axis labels are 1-based positions of SNPs (refer also to Supplementary Table 4).

We then characterized these 43 novel alleles and found that all novel alleles derive from core V genes we defined in a previous study according to their prevalence in antibody repertoires^14^ (Supplementary Figure 8A). This result further suggested that gene usage is critical in novel allele identification through Ig-seq dataset. Furthermore, the number of novel alleles did not correlate with the number of known alleles for a typical gene (Pearson correlation coefficient: 0.43) (Supplementary Figure 8B). However, IGHV1-69, the gene with the second largest known polymorphisms in IMGT, was found with up to 10 additional novel alleles. Since germline V gene polymorphisms have been implicated in immune response capability^1–3^, these novel alleles will facilitate the elucidation of the role of germline variants in disease susceptibility. Finally, we classified all identified SNPs (n=75) into two categories, replacement (R) SNPs and silent (S) SNPs, according to the variation of encoded amino acids. Overall, the R/S ratio for these SNPs was around 2 (1.88) (Supplementary Figure 8C). Nevertheless, the R/S ratio was larger for complementarity-determining region (CDR) SNPs (2.78) than framework region (FR) SNPs (1.41), which indicated a varied selection pressure between FRs and CDRs.

## Discussion

In this study, we comprehensively compared 5 NADTs with an emphasis on their performance in different scenarios. We identified 43 credible novel alleles through the filtration criteria informed by our benchmark results. We found that these NADTs possess a varied array of functionalities and distinct algorithms implemented in different languages (Table 1 and Supplementary Table 1). By exploiting a combination of *in silico* simulated and genuine Ig-seq datasets, we provided scenario-specific performance spectrums for these NADTs. As summarized in the Results section, both *TIgGER* and *IgDiscover* hit a greater balance between sensitivity and specificity in most scenarios than the other NADTs. In contrast, *LymAnalyzer* reported the greatest number of polymorphisms among NADTs, achieveing the highest sensitivity in all scenarios, however, at a great cost of specificity. *Partis* and *IMPre* were superior in overcoming challenges brought by SHMs and scarcity of minor alleles, respectively.

Counterintuitively, in our study *IgDiscover* rather than *TIgGER* exhibited higher efficiency in detecting novel alleles from SHM noise in GD-SHM. After careful examination, we identified the difference in candidate novel allele filtration between them. The quantitative filtration employed by *TIgGER* by default is mathematically more strict than the qualitative filtration used by *IgDiscover*. Combining the fact that a nonneglibigle fraction of unmutated sequences present also in IgG repertoires (Supplementary Figure 7), *IgDiscover* outperformed *TIgGER* even in SHM-enriched scenarios. However, ample sequencing depth is a prerequisite, because it favors the presence of enough unmutated sequence needed by *IgDiscover* to detect novel alleles. This is also true for all other NADTs because gene expression level was confirmed to be a general limitation for all NADTs in both simulated and genuine Ig-seq dataset (Table 3 and 4).

Although the unexpected observation above was not obtained by Ralph *et al.*^7^, they provided evidence that *TIgGER* can be completely compromised in handling with dataset of typically high SHM, which was possibly due to either the rarity of plurity sequence or the unqualified diversity of unmutated sequences. To maintain the originality of each NADT, we did not alter the suggested parameters and the results here may thus not represent the optimal performance for them. It is very likely that one can obtain greatly improved result when some key parameters are fine-tuned, a strategy that has been empolyed by Mikocziova *et al.*^19^. Despite the compromised sensitivity for *TIgGER* on particular datasets in this study, we agree with Ralph et al. that *IgDiscover* and *TIgGER* are more specific in novel allele detection than other NADTs, a major consideration of assigning more weight to them in the filtration scheme. We also noted some differences to Ralph *et al.*. The number of SNPs by which a novel allele departs from its nearest known allele (within a range from 1 to 3) are shown to exert negligible influence on *Partis*’s performance. However, our result revealed remarkable performance variance in detecting novel alleles separated from their nearest couterparts by SNPs of different number (i. e. 1 vs 3). This variance is probably caused by an error-prone procedure that *Partis* tries to manage – “comparing multiple hypotheses” (through which a complete set of individual SNPs contributing to a novel allele is determined). Noteworthy is that the step of initial removal of less-likely alleles in some cases can worsen the detection task for *Partis* because it can remove those less-likely but *bona fide* novel alleles that appears to harbor more than one SNPs.

Given all these findings, we suggest future studies to exploit strengths of different NADTs and present novel allele calls based on the consensus of more than one NADTs whenever genomic validation is unavailable, since none of the NADTs excels in all scenarios.

It should be noted that we considered only a single variable at a time. However, in real-world scenarios, a mixture of challenges represented by these studied variables coexiste and thus further complicate novel allele detection tasks. Moreover, we left polymorphisms of nucleotide insertion and deletion (INDEL) unaddressed because the algorithms empolyed by some NADTs are intrinsically incapable of capturing them (Supplementary Table 1). Nevertheless, INDEL can’t be neglected, especially in species whose germline sets are far from complete. In such cases, *IgDiscover* and *IMPre* are the only choices currently. Finaly, this study only focused on evaluation of NADTs’ performance based on antibody heavy chain repertoire datasets. Their efficiency with light chain and TCR repertoire datasets may vary due to differences inherent to these sequences (e. g. abence of SHM for TCR sequences).

Despite these limitations, our study based on a composite benchmark dataset provides insights into the performance of different NADTs and thus can guide bioinformaticians and immunologists in tool selection in future novel allele detection through these NADTs. Together with the flexible simulation tool and the novel alleles identified, our study may serve as a valuable reference and resource for immunoglobulin loci germline diversity researches as well as Ig-seq-based studies.

## Materials and Methods

### Samples from human subjects

A total of 28 samples from peripheral blood, tumor and normal tissues, and bone marrow were collected. Of these, 7 peripheral blood samples were derived from healthy individuals (without recent infection events), 6 peripheral blood samples were from hepatitis B virus-infected patients, 1 bone marrow sample and 2 peripheral blood samples were from graft-versus-host disease (GvHD) patients, 4 peripheral blood samples, 1 nomal intestine sample and 2 intestine tumor samples were from colorectal cancer (CRC) patients, 2 peripheral blood samples were from individuals involved in traffic accidents, and 3 peripheral blood samples were from patients with adolescent idiopathic scoliosis, sore throat, and chronic pharyngitis, respectively. Peripheral blood mononuclear cells (PBMCs) and bone marrow mononuclear cells were isolated using Ficoll (TBD Science) density-gradient centrifugation. The tissues were cut into small pieces and grind with liquid nitrogen. These experiments were handled under the guidelines of the Ethics Committee of Southern Medical University. For human naïve B cells isolation, PBMCs were counted and washed with DPBS supplemented with 1% bovine serum albumin (BSA), and then were stained with a cocktail of fluorescent conjugated antibodies, including ECD-CD19 (Beckman Coulter, A07770), FITC-IgD (Beckman Coulter, B30652), APC-CD27 (BD Bioscience, 561400), and 7-AAD (BD Bioscience, 559925). Human naïve B cells (CD19+IgD+CD27-7-AAD-) were sorted using a cell sorter (MoFlo XDP, Beckman Coulter) and collected for single-cell V(D)J sequencing.

### Library preparation and high-throughput sequencing

RNA purification was carried out using the RNeasy Mini Kit (Qiagen, 74106) according to the manufacturer’s instructions. Total RNA was used as a template to synthesize cDNA with a SMARTer RACE (Rapid Amplification of cDNA Ends) cDNA Amplification Kit (Clontech, 634928) according to the manufacturer’s protocol. Heavy chain variable regions were amplified using 1 μl of RT reaction product and 10 pmol of each primer in a 50 μl total reaction volume (KAPA HiFi HotStart ReadyMix, Roche) using the following thermal cycling program: 95 ℃ for 3 min; 30 cycles of 98 ℃ for 20 s, 60 ℃ for 15 s, and 72 ℃ for 15 s; 72 ℃ for 5 min. PCR products were purified using the Nucleospin Gel & PCR Clean-up kit (Macherey-Nagel, 704609.25) and subjected to library preparation using VAHTS Universal DNA Library Prep Kit (Vazyme, ND607-01). Libraries were quantified by capillary electrophoresis (Bio-Fragment analyzer, Bioptic). After quantification, libraries were pooled and sequenced on an Illumina platform (MiSeq PE300). All primers are listed in Supplementary Table 6.

### 10X Genomics single cell processing and next generation sequencing

The concentration of the single cell suspension was counted and adjusted to 1000 cells/μl. The single cell suspensions were loaded onto the Chromium Controller microfluidics device (10X Genomics) and processed using Chromium Next GEM Single Cell 5’ Kits v2 according to manufacturer’s protocol. The remaining procedures, including library construction, were performed according to the protocols of the Chromium Single Cell Human BCR Amplification Kit (10X Genomics). Following library construction, the BCR libraries were sequenced on an Illumina platform (NovaSeq 6000) using 2×150bp kit.

### Customization of reference sequences with artificially ‘novel’ V alleles

In this study, the germline reference sequences for V, D, and J genes were obtained from IMGT GENE-DB and provided as Supplementary Table 7. The artificially “novel” alleles for V genes were created for both simulated dataset and real Ig-seq dataset. Only germline reference sequences used in the simulation were extracted to serve as the initial reference sequences for the simulated dataset. The set of alleles subject to the artificial SNP generation for each dataset was selected according to the criteria defined as Table 2. We randomly created SNPs in the sequence of selected alleles. These artificial SNPs were set to locate in the first 280 bp of V genes at the 5’ ends to avoid the possible failure in novel allele detection caused by junctional modification. A pitfall here is that there exists a possibility that the rearranged sequences fail to be best aligned against the artificially novel sequences, and this brings challenges in the evaluation of novel allele identification for NADTs. Therefore, we performed pairwise alignment between customized reference sequences and the germline sequences contained in each dataset and removed those unaltered allele sequences that were found to be more similar to the germline sequences than the “novel” allele sequences. The novel alleles identified by NADTs were in fact the real-world germline sequences, while “novel” is just a concept relative to the altered germline reference sequences.

### Pipeline and parameters employed by 5 NADTs

The pair-end simulated dataset and bulk sequencing dataset were firstly assembled using *PEAR* (v0.9.6). The successfully assembled sequences were then taken as the input for *IgDiscover* and *LymAnalyzer*. As *TIgGER* can only accept a formatted database of well-annotated sequences as input, we further annotated and formatted the assembled sequences with *IgBLAST* (v2.8.0+) and *Change-O* toolkits (v0.4.4), respectively (*IgBLAST* was selected for its excellent performance^20^ and easy output format conversion through *Change-O* toolkits). For *IMPre* and *Partis*, the input assembled sequences were corrected in a forward orientation at first. The script employed by *IMPre* (‘*IMPre.pl*’) was modified to enable germline reference customization. The revised script, ‘*IMPre_revised.pl*’, can be found on the Github (https://github.com/Xiujia-Yang/IMPlAntS). All parameters used by the five NADTs were set in default or as suggested. We provided the detailed commandline parameters as below,

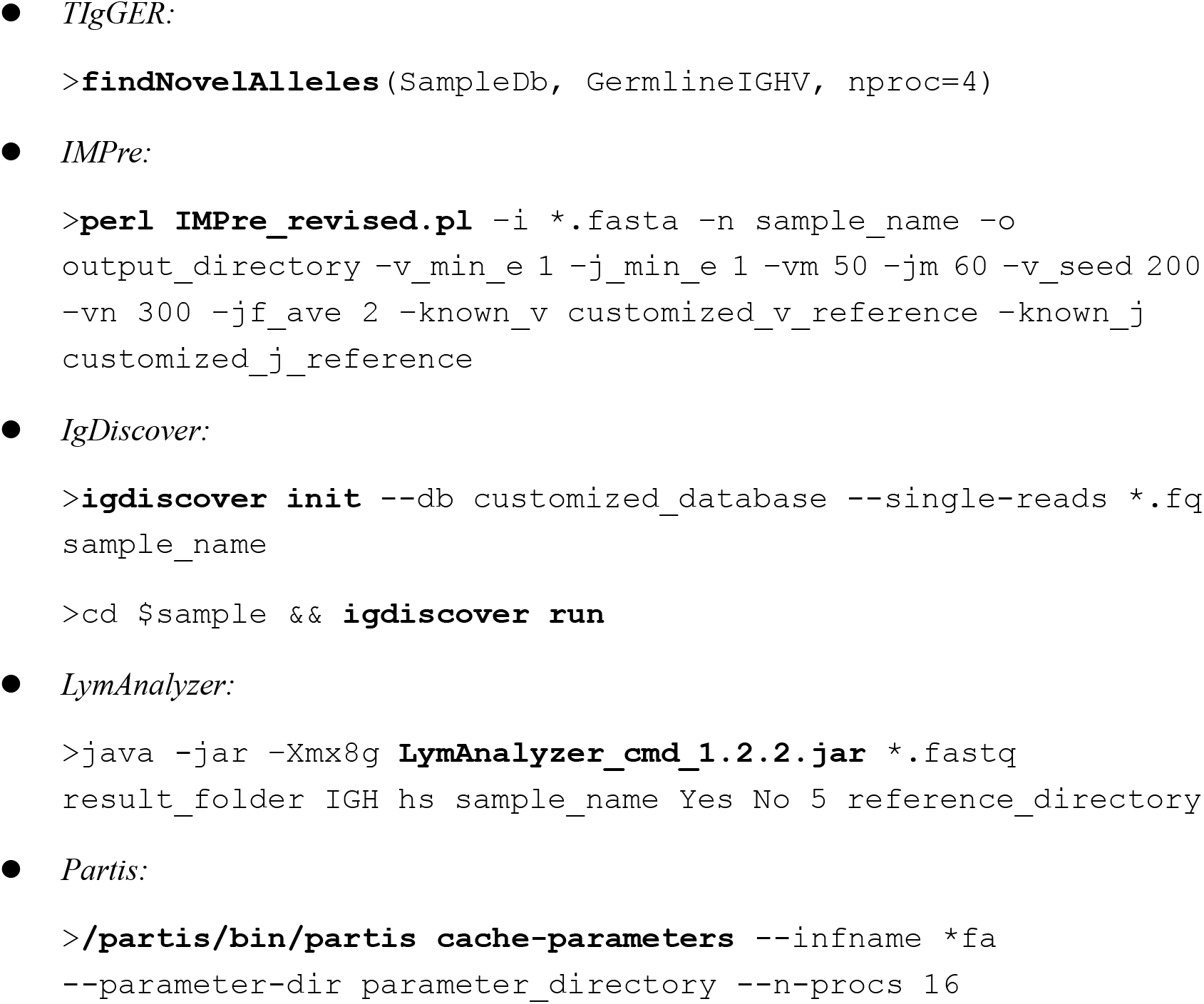

### Sensitivity and specificity calculation

Sensitivity is defined as the proportion of true positives that are correctly identified among all true positives, whereas specificity is defined as the proportion of true positives among all the identified positives. For individual SNPs (SNP level), a hit is considered as a true positive only when its nearest allele (same as the allele selected for artificial SNP generation), loci and nucleotide variant are correct at the same time. For individual sequences (allele level), a hit is considered a true positive only when it covers all the genuine SNPs and contains no mismatches with the genuine novel sequence in all other reported loci. A schematic diagram is provided here to demonstrate the cases of true positive and false positive in identifying individual sequences for novel alleles.

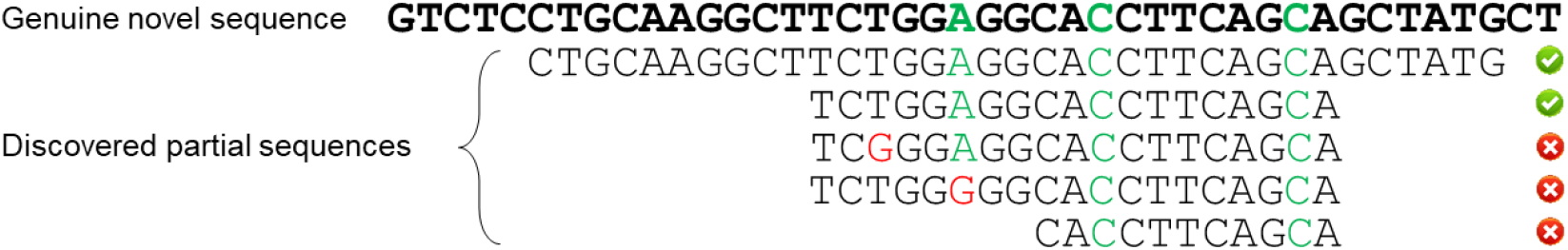

### Schematic diagram of true positive and false positive of discovered novel sequences

<u>The top sequence in bold represents the genuine novel sequence while the bottom sequences represent the partial/full-length sequences discovered by NADTs. The nucleotides marked in green represent the genuine SNPs while those in red are mismatches with the genuine novel sequence either in SNP loci or non-SNP loci. An identified sequence is accepted as a true positive only when it covers all the genuine SNPs and contains no mismatch with the genuine novel sequence in all other loci.</u>

### Germline V allele identification through single naïve B cell sequencing dataset

*Cell Ranger* (v3.1.0) was exploited to preprocess the raw single naïve B cell sequencing dataset. Contig assembly, annotation, and clonotype analysis were performed using “cellranger vdj” with the *Cell Ranger* V(D)J compatible reference (refdata-cellranger-vdj-GRCh38-alts-ensembl-3.1.0). Then the assembled *contig* sequences (“all_contig.fasta”) of the two replicates for each donor were pooled and then annotated using *IgBLAST* (v2.8.0+) with the germline references obtained from IMGT/GENE-DB (refer to above). Afterwards, the V segment (or allele) sequence was extracted from each annotated sequence and then each unique V segment sequences was counted. It is worth mentioning here that those short V segment sequences were merged into the longer ones provided they are with the same V allele annotation as the longer ones and were included in them. The counts for the short V segment sequences were also added to the longer ones. We discarded those with a length less than 290 bp or with a count less than 10 and determined the most frequent V segment sequence for each gene as the most confidential germline sequence for a gene. Apart from that, we also retained the second most frequent V segment sequence for a gene provided that its abundance was at least one tenth of that of the most frequent one.

### Ig-seq dataset classification criteria

All enrolled Ig-seq datasets (i.e. 424 RACE datasets and 263 multiplex datasets mentioned in the Results section) were analyzed using MiXCR (v3.0.7) per the method in our previous study^14^. After clonotype assembly, a constant gene will be assigned for each clone if antibody sequences from this clone cover constant region. The isotype (i.e. IgM, IgD, IgG, IgA, and IgE) was extracted for each clone and the clone-level isotype frequency was calculated for each dataset. IgM and IgD are deemed as SHM-sparse isotypes while IgG, IgA and IgE are deemed as SHM-enrich isotypes^21^. Datasets will be classified as IgM datasets if they contain more SHM-sparse isotypes than SHM-enrich isotypes, otherwise IgG datasets. Constant genes were required to be assigned for more than a half number of clones in each dataset. All 687 Ig-seq datasets we enrolled in this study met this requirement.

### V allele sequences from public databases and independent reports

To double validate novel alleles we identified through NADTs, we collected antibody heavy chain V allele sequences from four public databases (IMGT/GENE-DB, http://www.imgt.org/genedb/; IgPdb, https://cgi.cse.unsw.edu.au/~ihmmune/IgPdb/information.php; VBASE2^22^, http://www.vbase2.org/; Lym1K^23^, http://maths.nuigalway.ie/biocluster/database/) and nine independent reports^5,6,19,24–29^ and compared them with identified novel alleles. Before the sequence comparison, degenerate bases or N nucleotides in collected allele sequences were substituted with ‘A’, ‘C’, ‘G’, or ‘T’, accordingly. Novel alleles whose sequences were identical to any of the sequences from a source were considered validated novel alleles. As the set of V allele sequences used as germline reference to identify novel alleles is not as complete as those in the later release of IMGT/GENE-DB, several novel allele sequences were included in the later release of IMGT/GENE-DB and thus were validated in it. All collected V allele sequences are outlined in Supplementary Table 8.

## Data availability

In-house data including paired single naïve B cell and bulk sequencing dataset and unpaired bulk sequencing dataset is stored in NCBI SRA database under accession number PRJNA732986.

## Acknowledgement

This study was supported by the National Natural Science Foundation of China (NSFC) (31771479 to Zhenhai Zhang), NSFC Projects of International Cooperation and Exchanges of NSFC (61661146004 to Zhenhai Zhang), the Local Innovative and Research Teams Project of Guangdong Pearl River Talents Program (2017BT01S131 to Zhenhai Zhang) and Guangdong-Hong Kong-Macao-Joint Labs Program from Guangdong Science and Technology (2019B121205005 to Xueqing Yu).

## Author contributions

X. Y., Y. Z., H. Z., S. C., and C. L. performed bioinformatics analyses on the data, Q. W. and J. G. collected samples and conducted the biological experiments. X. Y., X. Y. and Z. Z. wrote the manuscript. Z. Z. conceived the project.

## References

1. Lingwood, D. et al. Structural and genetic basis for development of broadly neutralizing influenza antibodies. Nature. 489, 566–570 (2012).

2. Avnir, Y. et al. IGHV1-69 polymorphism modulates anti-influenza antibody repertoires, correlates with IGHV utilization shifts and varies by ethnicity. Sci. Rep. 6, (2016).

3. Parks, T. et al. Association between a common immunoglobulin heavy chain allele and rheumatic heart disease risk in Oceania. Nat. Commun. 8, (2017).

4. Lees, W. et al. OGRDB: a reference database of inferred immune receptor genes. Nucleic Acids Res. 48, D964–D970 (2020).

5. Corcoran, M. M. et al. Production of individualized V gene databases reveals high levels of immunoglobulin genetic diversity. Nat. Commun. 7, (2016).

6. Gadala-Maria, D., Yaari, G., Uduman, M. & Kleinstein, S. H. Automated analysis of high-throughput B-cell sequencing data reveals a high frequency of novel immunoglobulin V gene segment alleles. Proceedings of the National Academy of Sciences. 112, E862–E870 (2015).

7. Ralph, D. K. & Matsen, F. A. Per-sample immunoglobulin germline inference from B cell receptor deep sequencing data. PLOS Computational Biology. 15, e1007133 (2019).

8. Yu, Y., Ceredig, R. & Seoighe, C. LymAnalyzer: a tool for comprehensive analysis of next generation sequencing data of T cell receptors and immunoglobulins. Nucleic Acids Res. 44, e31 (2016).

9. Zhang, W. et al. IMPre: An Accurate and Efficient Software for Prediction of T- and B-Cell Receptor Germline Genes and Alleles from Rearranged Repertoire Data. Frontiers in Immunology. 7, (2016).

10. Marcou, Q., Mora, T. & Walczak, A. M. High-throughput immune repertoire analysis with IGoR. Nat. Commun. 9, (2018).

11. Safonova, Y., Lapidus, A. & Lill, J. IgSimulator: a versatile immunosequencing simulator. Bioinformatics. 31, 3213–3215 (2015).

12. Weber, C. R. et al. immuneSIM: tunable multi-feature simulation of B- and T-cell receptor repertoires for immunoinformatics benchmarking. Bioinformatics. 36, 3594–3596 (2020).

13. Yermanos, A. et al. Comparison of methods for phylogenetic B-cell lineage inference using time-resolved antibody repertoire simulations (AbSim). Bioinformatics. 33, 3938–3946 (2017).

14. Yang, X. et al. Large-scale analysis of 2,152 Ig-seq datasets reveals key features of B cell biology and the antibody repertoire. Cell Rep. 35, 109110 (2021).

15. Huang, W., Li, L., Myers, J. R. & Marth, G. T. ART: a next-generation sequencing read simulator. Bioinformatics. 28, 593–594 (2012).

16. Watson, C. T. et al. Complete Haplotype Sequence of the Human Immunoglobulin Heavy-Chain Variable, Diversity, and Joining Genes and Characterization of Allelic and Copy-Number Variation. The American Journal of Human Genetics. 92, 530–546 (2013).

17. Budeus, B. et al. Complexity of the human memory B-cell compartment is determined by the versatility of clonal diversification in germinal centers. Proceedings of the National Academy of Sciences. 112, E5281–E5289 (2015).

18. Ghraichy, M. et al. Maturation of the Human Immunoglobulin Heavy Chain Repertoire With Age. Frontiers in Immunology. 11, (2020).

19. Mikocziova, I. et al. Polymorphisms in human immunoglobulin heavy chain variable genes and their upstream regions. Nucleic Acids Res. 48, 5499–5510 (2020).

20. Zhang, Y. et al. Tools for fundamental analysis functions of TCR repertoires: a systematic comparison. Brief. Bioinform. 21, 1706–1716 (2020).

21. Kitaura, K. et al. Different Somatic Hypermutation Levels among Antibody Subclasses Disclosed by a New Next-Generation Sequencing-Based Antibody Repertoire Analysis. Frontiers in Immunology. 8, (2017).

22. Retter, I. VBASE2, an integrative V gene database. Nucleic Acids Res. 33, D671–D674 (2004).

23. Yu, Y., Ceredig, R. & Seoighe, C. A Database of Human Immune Receptor Alleles Recovered from Population Sequencing Data. J. Immunol. 198, 2202–2210 (2017).

24. Gadala-Maria, D. et al. Identification of Subject-Specific Immunoglobulin Alleles From Expressed Repertoire Sequencing Data. Frontiers in Immunology. 10, (2019).

25. Gidoni, M. et al. Mosaic deletion patterns of the human antibody heavy chain gene locus shown by Bayesian haplotyping. Nat. Commun. 10, (2019).

26. Thörnqvist, L. & Ohlin, M. Critical steps for computational inference of the 3′-end of novel alleles of immunoglobulin heavy chain variable genes - illustrated by an allele of IGHV3-7. Mol. Immunol. 103, 1–6 (2018).

27. Vázquez Bernat, N. et al. High-Quality Library Preparation for NGS-Based Immunoglobulin Germline Gene Inference and Repertoire Expression Analysis. Frontiers in Immunology. 10, (2019).

28. Wang, Y. et al. Genomic screening by 454 pyrosequencing identifies a new human IGHV gene and sixteen other new IGHV allelic variants. Immunogenetics. 63, 259–265 (2011).

29. Wendel, B. S., He, C., Crompton, P. D., Pierce, S. K. & Jiang, N. A Streamlined Approach to Antibody Novel Germline Allele Prediction and Validation. Frontiers in Immunology. 8, (2017).

